# Fetal Twin: a mechanistic computational model of fetal physiology for heart-rate-variability biomarker research

**DOI:** 10.64898/2026.07.14.738362

**Authors:** Martin G. Frasch

## Abstract

Fetal-monitoring biomarkers for neonatal hypoxic-ischemic brain injury face a structural gap: the mechanistic ground truth that would label a training set — perfusion pressure, the moment of decompensation, the injury time course — cannot be measured at scale or ethically in human pregnancy or labor, and generative synthetic data carry no mechanistic labels. We address this with a mechanistic computational model of the fetal cardiovascular, autonomic, and metabolic response to controlled hypoxic stress, and use it to test how beat detection and acquisition fidelity alter the interpretation of fetal-heart-rate-variability (HRV) biomarkers. The model integrates these systems forward in time across antepartum development (gestational-age growth scaling) and intrapartum stress (umbilical-cord occlusions), emitting synthetic monitoring signals (fetal heart rate, RR intervals) co-registered with model-computed latent labels (pH, base deficit, lactate, perfusion pressure, decompensation and injury states). In a fetal-sheep-derived autonomic-loop configuration we report three results. First, a phase-accumulator beat detector shows that the apparently physiologic baseline HRV of an earlier build was largely a detector artifact, and a noise-off control shows beat-to-beat variability requires an explicit stochastic driver rather than self-sustained autonomic oscillation. Second, a sampling-fidelity sweep yields a fidelity-matched selection rule: a deceleration-area biomarker is preserved at CTG-grade 4 Hz whereas RMSSD is corrupted there (inflated about 8-fold by timing quantization) and recovers only at fetal-ECG rates. Third, autonomic modulation alone does not reproduce the published RMSSD rise-then-collapse — a negative result that motivates, but does not prove, an intrinsic sinoatrial-pacemaker contribution as a testable hypothesis. This is an in-silico, hypothesis-generating study: the model is not validated for individual fetal prediction, clinical risk estimation, or clinical decision-making. The model is implemented as **Fetal Twin** (engine `fetaltwin`), a source-available research instrument released under a noncommercial license, together with all figure configurations, so that these controlled experiments are reproducible.

**Key Points:**

- Progress on fetal-monitoring biomarkers for neonatal brain-injury risk is constrained by a structural gap: the mechanistic ground truth that would label a training set — perfusion pressure, the moment of cardiovascular decompensation, the time course of injury — cannot be measured at scale or ethically during human pregnancy or labor.
- We present **Fetal Twin** (source-available engine `fetaltwin`), a publicly available, noncommercially-licensed mechanistic testbed for fetal physiological development. It integrates the fetal cardiovascular, metabolic, and autonomic systems forward in time across **antepartum development** (gestational-age growth scaling) and **intrapartum stress** (umbilical-cord occlusions), and emits synthetic monitoring signals (fetal heart rate, RR intervals) co-registered with model-computed latent labels (pH, lactate, perfusion pressure, decompensation and injury states).
- The name’s digital-twin connotation is deliberate but bounded: Fetal Twin is a **mechanistic, population-level twin of fetal physiology used as a research instrument** — not a validated, patient-specific clinical digital twin or predictor. Its purpose is to interrogate what candidate biomarkers can and cannot mean, via in-silico controls impossible in vivo — swapping the beat detector, turning a noise source off, ablating a reflex, or quantizing the signal to a monitor’s sampling grid.
- Demonstrations in a fetal-sheep-derived autonomic-loop configuration show that beat-to-beat HRV *amplitude* can be a numerical artifact of the beat detector, and that in this model class beat-to-beat variability requires an explicit stochastic driver — it does not arise as a self-sustained oscillation of the deterministic autonomic loop.
- A further demonstration establishes a **fidelity-matched biomarker-selection rule** — a deceleration-area biomarker survives CTG-grade 4 Hz sampling whereas RMSSD is corrupted at that rate and needs fetal-ECG timing — and a negative result shows the published RMSSD “rise-then-collapse” is not reproducible from autonomic modulation in this model class, motivating (but not proving) an intrinsic-pacemaker hypothesis.

## 1 Introduction

Continuous FHR monitoring — by cardiotocography (CTG) using Doppler ultrasound, or by fetal electrocardiography (fetal ECG) — is the primary tool for intrapartum surveillance, yet its positive predictive value for the clinically important outcome is poor. That outcome is neonatal hypoxic-ischemic brain injury, driven proximately by fetal cardiovascular decompensation (systemic hypotension) during labor, not by fetal acidemia per se [2, 4, 3].

Two structural obstacles have impeded better biomarkers. First, **absence of ground-truth labels**: continuous mechanistic ground truth — arterial and cerebral perfusion pressure, the moment of decompensation, the time course of injury — cannot be measured at scale or ethically during human pregnancy or labor, so candidate predictors have no ground-truth training signal. Generative synthetic-data methods (generative adversarial networks; kernel-density / morphological resampling) are effective at reproducing the marginal and temporal statistics of real recordings, and are the appropriate tool when the goal is data augmentation; but because they model the signal rather than the physiology that produced it, they cannot supply mechanistic labels. Second, even where a biomarker is promising, it is rarely clear **what physiological quantity it actually reflects**, or whether its apparent signal is a property of the acquisition and analysis pipeline rather than of the fetus.

A mechanistic simulator addresses both. Prior mechanistic models of the fetal circulation have reproduced the shape of variable decelerations from cord-compression physiology [7] and the allometric scaling of the fetal circulation across gestation [8], but were not built to emit ground-truth-labeled training signals for biomarker evaluation. If a model integrates the fetal cardiovascular, metabolic, and autonomic dynamics forward in time — across antepartum development and intrapartum stress — every latent quantity is available as a noise-free, continuously-sampled label co-registered with the synthetic monitoring signal. The model also permits experiments impossible in vivo — turning a noise source off, ablating a reflex pathway, or quantizing the signal to a target monitor’s sampling grid — that reveal what a candidate biomarker is really measuring.

Here we describe **Fetal Twin**, a source-available, noncommercially-licensed testbed for fetal physiological development, implemented in the `fetaltwin` engine as an extension of the Wang et al. (2015) fetal umbilical-cord-occlusion (UCO) model [1] with gestational growth scaling after van Willigen et al. (2024) [8]. We state our claims as an explicit hierarchy, strongest first, so that the weaker ones cannot undermine the stronger:

1. **Primary (a mechanistic model and the controlled experiments it makes possible)**. A ground-truth-labeled mechanistic model spanning antepartum and intrapartum fetal physiology, together with the in-silico controls — swapping the beat detector, disabling a noise source, ablating a reflex, quantizing to a monitor’s sampling grid — that make it possible to interrogate what a monitoring biomarker actually measures. The source-available engine is provided so that every experiment below is reproducible.
2. **Secondary (the within-model methodological findings)**. In a fetal-sheep-derived autonomic-loop configuration, certain HRV behaviors are artifacts of the observation pipeline or are injected features unless explicitly modeled otherwise, and biomarker information content depends on acquisition fidelity in a quantifiable way. These hold for this model class and follow from the controls the model makes possible.
3. **Tertiary (a hypothesis it generates)**. The failure of autonomic modulation to reproduce the published “rise-then-collapse” HRV trajectory suggests, but does not establish, an intrinsic-pacemaker contribution.

We do not claim a validated clinical predictor; the model is calibration-grade, and we are explicit throughout about what it must be valid for and what it is not (Section 4).

## 2 Methods

### 2.1 The mechanistic model

The simulator implements and extends the combined fetal-sheep model of Wang et al. (2015) [1]. The circulation, oxygen/CO_2_ transport, and baro/chemoreflex structure are **inherited from Wang 2015**; the acid–base/lactate coupling, the autonomic HRV mechanisms, the intrinsic-HRV surrogate, the physiological deceleration pathway, the measurement-fidelity model, and the gestational growth scaling are **added in this work** (parameter deltas from Wang 2015 are enumerated in the released configuration and the Supplementary Methods). The coupled ODE system integrates **33 states** spanning four subsystems:

- **Combined circulation** — arterial, venous, cerebral, and umbilical compartments and a time-varying-elastance ventricular pump yielding instantaneous arterial pressure and mean arterial pressure (MABP).
- **Oxygen / CO**_***2***_ **/ acid-base** — compartmental O_2_ and CO_2_ content, lactate, an oxygen-gated anaerobic glycolytic pathway (GL → 2*·*LA + 2·H^+^), an explicit bicarbonate buffer, pH by Henderson–Hasselbalch, and base deficit by the Stewart approach. An O_2_-gated, acidosis-suppressed oxidative lactate-clearance sink couples lactate to the acid-base state so that high lactate co-occurs with acidemia.
- **Autonomic control** — baro- and peripheral-chemoreflex afferents and vagal / sympathetic efferents; the instantaneous heart period is decomposed as T = T_0_ + ΔT_s + ΔT_va (intrinsic, sympathetic, vagal contributions).
- **Intrinsic-HRV surrogate** — a per-beat multiplicative modulation of the regulated period, T_final = T_eff *·* (1 + g *δ*_k), with *δ*_k a second-order autoregressive (AR(2)) process indexed by beat number; g = 0 recovers deterministic timing, so intrinsic variability and autonomic modulation are separable. The AR(2) form is the minimal linear process able to reproduce a damped-oscillatory beat-to-beat autocorrelation (a broadband peak rather than a pure Lorentzian), and its coefficients were distilled to match the short-lag autocorrelation of a separate stochastic sinoatrial coupled-clock pacemaker model. We stress that this surrogate is a **phenomenological placeholder for intrinsic pacemaker variability, not a mechanistic sinoatrial-node model**; statements below about an “intrinsic” origin are therefore provisional and motivate, rather than instantiate, a mechanistic pacemaker.

#### Developmental staging

The model spans two stages. *Antepartum:* a gestational growth-scaling transformation after van Willigen et al. (2024) [8] applies allometric laws (verified against the source) to scale term-fetus hemodynamics to any gestational age over ~20–40 weeks; no-occlusion baselines at 28/34/40-week equivalents are integrator-stable and reproduce the expected fall in cardiac output toward preterm. *Intrapartum:* UCO forcing is a deterministic progression of occlusion frequency, duration, and degree (e.g., a 1:2.5 occlusion-to-gap cadence), with bounded per-event variability. The demonstrations in Section 3 use the intrapartum stage on the term (sheep-calibrated) configuration.

#### Physiological decelerations (V3.0)

A complete UCO produces a physiological fetal deceleration (resting FHR ~140→ nadir ~75–85 bpm) driven by a flow-weighted arterial-O_2_ peripheral chemoreflex, grounded in the Lear & Gunn fetal-sheep literature [2]. (An earlier formulation produced a paradoxical tachycardia; the diagnostic that located and corrected this is reported in the repository.)

### 2.2 Numerics and reproducibility

The system is integrated with a BDF stiff solver. Two interchangeable right-hand-side backends — a pure-Python reference and a Numba-JIT backend (~5× faster) — are validated pointwise-equivalent to rtol = 1e-9. Scenarios are specified as YAML; outputs are CSV time series plus beat-to-beat RR and (optionally) 100 Hz pulsatile-MABP. A browser-based interactive interface (a Streamlit application) is also released for building umbilical-cord-occlusion scenarios, running the engine, and replaying a simulation as a scrolling cardiotocography strip, lowering the barrier to exploring the testbed (Section 7). All code, configuration for the experiments below, and the analysis scripts are released (Section 7).

### 2.3 Beat detection

Beat (R-peak) times are detected from a **cardiac-phase accumulator** state *φ* that integrates d*φ*/dt = 1/T_period and emits a beat at each integer crossing of *φ*. This is robust to a time-varying period. We contrast it (Section 3.1) with a naive detector that reconstructs phase as (t mod T) − T/2, which is *not*.

### 2.4 HRV mechanisms

The vagal heart-period drive is split into a slow tonic pathway (`tau_T_va = 6 s`, integrated baro/chemoreflex tone) and a fast phasic pathway (`tau_T_va_fast = 0.4` s) that carries an Ornstein–Uhlenbeck vagal-noise process at near beat-to-beat scale. An optional episodic fetal-breathing module adds respiratory-sinus-arrhythmia modulation. A vagal-exhaustion state *θ* applies a (1 − *θ*) gain to phasic HRV, so accumulating exhaustion suppresses variability.

### 2.5 Measurement-fidelity model

To emulate a monitor of sampling rate f_s, beat times are quantized to the acquisition grid, 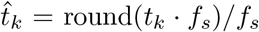, and RR intervals recomputed as 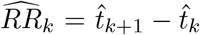, while a morphological area biomarker is computed on the FHR resampled to the same grid. CTG Doppler corresponds to f_s ≈ 4 Hz (*±*0.25 s per-beat timing jitter); fetal ECG to beat-resolved timing [5]. The sweep is run over f_s ∈ {4, 8, 16, 32, 128, 250} Hz (released analysis; Figure 4).

### 2.6 Ground-truth labels and predictor scoring

For each time step the simulator records the synthetic signal with the latent state as labels: perfusion pressures; pH / base deficit / lactate; a cardiovascular-decompensation reserve; a sustained-perfusion envelope; a cerebral injury burden; and a myocardial-reserve state. As an illustrative predictor we score the Roux et al. (2021) Distance-to-Healthy (DTH) metric [3] and time-domain HRV (RMSSD, SDNN) on the synthetic RR against the labels, by Spearman correlation and by lead time to a decompensation threshold.

## 3 Results

### 3.1 HRV amplitude in these models can be numerical

The correct instantaneous cardiac phase is the running time integral of the instantaneous rate, 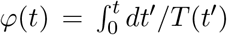, and a beat occurs at each unit increment of *φ*. A *naive* detector instead reconstructs phase algebraically from the current period, 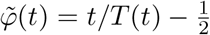, and emits a beat at each downward zero-crossing of the resulting sawtooth. When T is constant the two agree; when T varies, differentiating the naive form gives 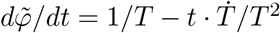, i.e. a spurious term 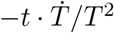 whose magnitude grows with elapsed time t. Thousands of cycles into a long run this term becomes comparable to the true 1/T rate, so a slow drift in T injects extra zero-crossings — spurious beats. We quantified this by counting detected beats with no true beat (from the phase-accumulator reference) within 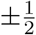 of the local mean RR: a 100-min UCO run came out **94.5% spurious beats**. With correct phase-accumulator detection the baseline RMSSD that an earlier implementation had targeted as physiologic (~8.5 ms) collapsed to ≈ **0.04 ms** — the prior amplitude had been the detector’s own jitter, and a single 6 s vagal effector was low-passing the injected noise away before it could reach beat-to-beat scale. Restoring physiologic variability required the explicit fast/slow vagal split (Section 2.4) and recalibrating the vagal-noise amplitude, after which baseline RMSSD ≈ **9.5 ms**, SDNN ≈ 18.7 ms at ~120 bpm, with the pure-Python and Numba backends agreeing pointwise (Figure 1).

**Figure 1.**
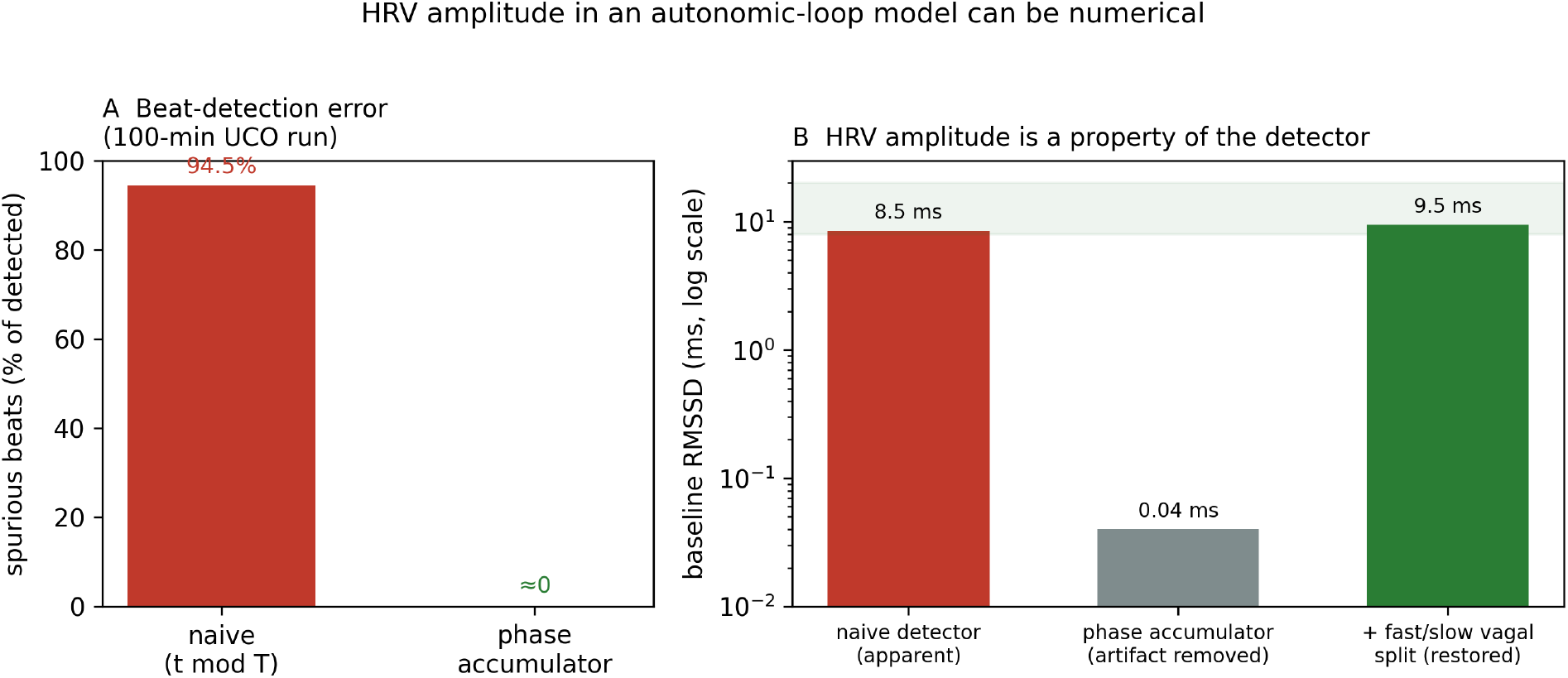
HRV amplitude in an autonomic-loop model can be numerical. (A) A naive beat detector that reconstructs cardiac phase as (t mod T) from a time-varying period produces 94.5% spurious beats on a 100-min UCO run; the phase-accumulator detector produces essentially none. (B) Baseline RMSSD an earlier build had targeted as physiologic (~8.5 ms) collapses to ~0.04 ms under correct phase-accumulator detection — the amplitude was the detector’s own jitter — and is restored to ~9.5 ms (physiologic band shaded) only after the explicit fast/slow vagal split and vagal-noise recalibration (log scale). Values from the documented reanalysis (Results 3.1).

The 94.5% figure is the historical measurement that motivated replacing the naive event with the phase accumulator; it is reported as the observation on the affected run, not as a controlled parameter sweep. The general lesson does not depend on the exact number: whenever beats are *detected* from an integrated period rather than *emitted* as intrinsic events, a time-varying period makes the reconstruction ill-posed — a class of failure that a pacemaker-native beat generator would avoid by construction (Section 4).

*Methodological takeaway:* in simulators where HRV is injected through a low-pass autonomic effector, reported beat-to-beat HRV can be dominated by detector and filter artifacts. Amplitude alone is not evidence of physiology; the generating mechanism and the detector must be inspected.

### 3.2 In this model class, beat-to-beat variability requires an explicit stochastic driver

We use “emergent” in a specific operational sense: variability is emergent if the *deterministic* system (all explicit noise sources removed) produces sustained beat-to-beat fluctuation of its own accord for example a self-sustained loop oscillation. By that test the variability here is not emergent. With the fully deterministic system — both the injected vagal noise (*σ* = 0) and the intrinsic-HRV surrogate off — the loop carries only **0.08%** of the stochastic run’s spectral power: the autonomic baroreflex loop does not self-oscillate, and there is no spontaneous Mayer-wave-type generator. Beat-to-beat variability in this implementation therefore *requires an explicit stochastic driver*; the loop modulates the variability it transmits but does not itself produce substantial spontaneous variability in its absence (Figure 2).

**Figure 2.**
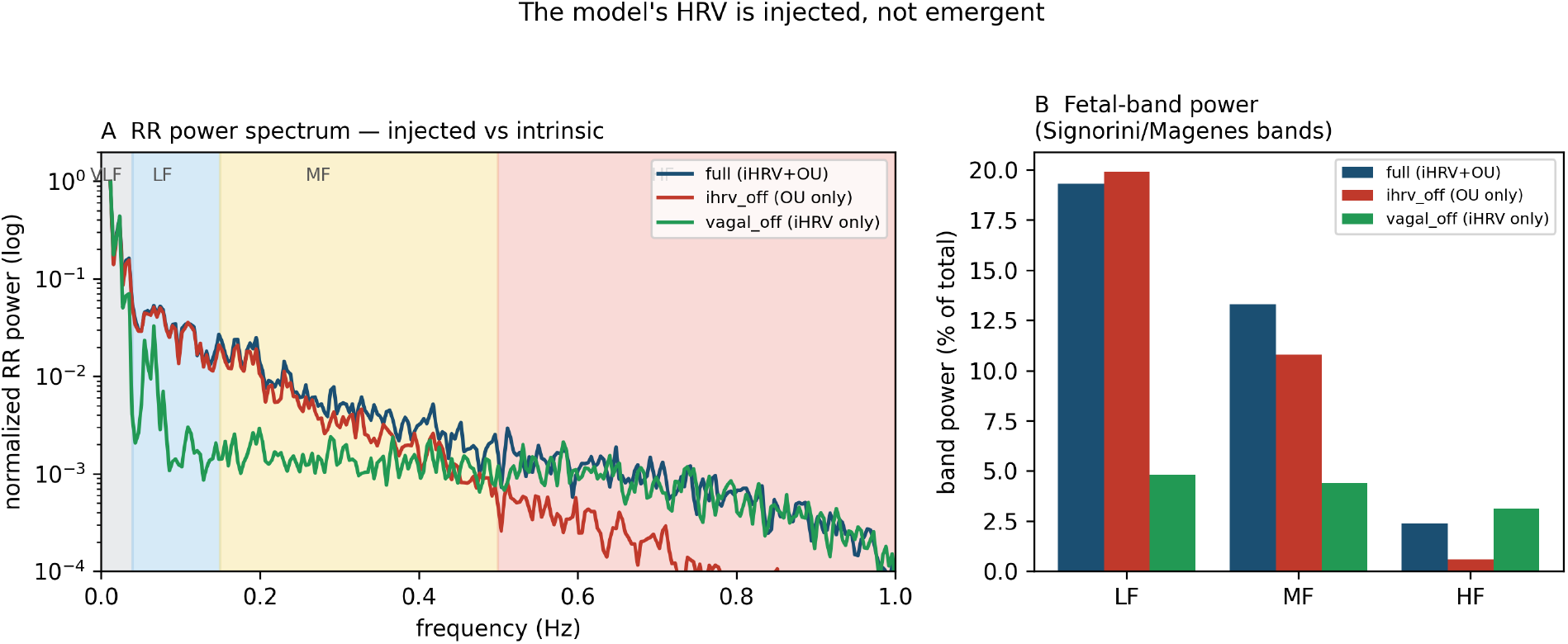
Beat-to-beat variability requires an explicit stochastic driver; it does not arise as a self-sustained oscillation of the deterministic loop. (A) Normalized RR power spectra for the full model (intrinsic surrogate + injected Ornstein–Uhlenbeck vagal noise), OU-noise-only, and intrinsic-surrogate-only, with the fetal Signorini/Magenes bands shaded. (B) Band power as a percentage of total. Removing the injected OU noise collapses the LF and MF bands (19% → 4.8%, 13% → 4.4%), showing the low-/mid-frequency structure is carried by the injected process rather than generated by the loop; the fetal HF band is near-empty (2.4%) in all cases because the model has no fetal-breathing generator. 30-min stationary baseline run.

The band decomposition localizes the driver (Figure 2B; percentages are of total power, using the Signorini/Magenes fetal-band frequency ranges [6] as descriptive bins — we do not claim these bands map one-to-one onto distinct physiological generators). With the injected Ornstein–Uhlenbeck vagal noise present the spectrum is VLF/LF/MF-dominated (VLF 63%, LF 19%, MF 13%); removing that noise so only the intrinsic surrogate remains **collapses the LF and MF bands (to 4.8% and 4.4%)**, showing the low-/mid-frequency structure is dominated by the imposed stochastic process rather than generated endogenously by loop dynamics. The high-frequency band (0.5–1.0 Hz) is essentially **empty (2.4%)** throughout — as expected, since the model has no fetal-breathing generator, the dominant fetal HF source; enabling the fetal-breathing module fills this band (to 49.4%) with a clean peak at the breathing frequency.

*Methodological takeaway:* spectral features attributed to autonomic loop dynamics in such models may instead reflect the imposed noise’s shape. Separating driver-dominated from endogenously-generated variability requires the noise-off control.

### 3.3 Which axis does an HRV predictor read — and does it still warn early?

On a 1:2.5 UCO decompensation run, sliding-window RMSSD correlates strongly with the vagal-exhaustion state (*ρ* = −0.51, p ≈ 1e-16) and with cerebral O_2_ and MABP, but is statistically independent of pH (*ρ* = −0.05, n.s.) and lactate. RMSSD thus reads the **autonomic/hypoxic-exhaustion axis**, not the acid-base axis directly — consistent with in-vivo evidence that fetal RMSSD tracks vagus nerve activity fluctuations [9].

Importantly, this does not make the HRV predictor useless for metabolic decompensation. On the current substrate (physiological decelerations, faithful lactate–acid coupling, intrinsic HRV on by default), the DTH predictor on synthetic RR rises in trend as pH falls (*ρ*(DTH, pH) = −0.43 with intrinsic HRV on; −0.65 with it off), and in this run its alarm fires roughly an hour before pH crosses 7.15 (alarm ≈ 10 min, pH < 7.15 ≈ 67 min; a lead of ~57 min). The autonomic trajectory the predictor tracks thus *precedes* the acid crossing here, so an autonomic-axis biomarker can still deliver early warning of a metabolic event. Intrinsic HRV slightly dilutes the correlation but does not move the alarm time — consistent with it being a decompensation-neutral overlay that enriches short-term HRV structure (Figure 3).

**Figure 3.**
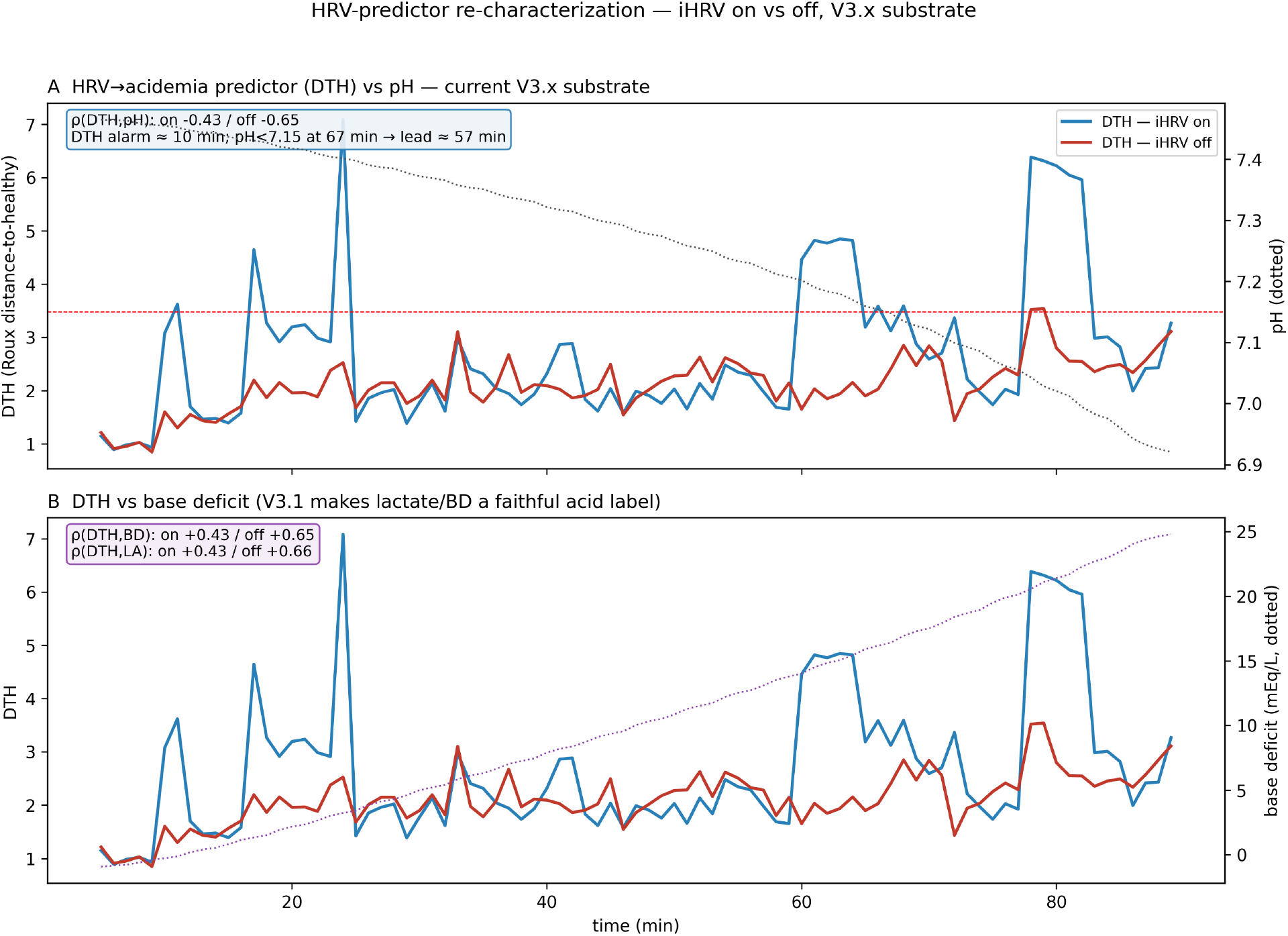
An HRV predictor reads the autonomic axis yet still warns early. Roux et al. (2021) Distance-to-Healthy (DTH) [3] scored on the synthetic RR over a 1:2.5 UCO decompensation run, with intrinsic HRV on (blue) and off (red). (A) DTH versus ground-truth pH (dotted): *ρ*(DTH, pH) = −0.43 (on) / −0.65 (off); the DTH alarm fires at ~10 min and pH < 7.15 at ~67 min, a ~57-min lead. (B) DTH versus base deficit — lactate and base deficit are faithful acid labels (*ρ* ≈ +0.43 / +0.65). Intrinsic HRV dilutes the correlation but does not move the alarm time.

Across five independent noise seeds at this severity, the lead is **invariant at 57 min** (IQR [57, 57]; alarm 10 min and pH < 7.15 crossing 67 min in every seed; 0/5 runs excluded by the metabolic-runaway guard), while the seed-dependent HRV noise moves the correlation — *ρ*(DTH, pH) ranges −0.34 to −0.78 (median −0.49) — but not the alarm or the lead (Figure S1; Supplementary Methods S8). The lead is stable because both endpoints are set by the near-deterministic acid-base trajectory and an early, reliable DTH alarm, whereas *ρ* reflects the stochastic short-term HRV structure. Two caveats remain, and we are explicit about them. First, the lead is **regime-dependent**: a severity sweep (3 seeds each) shows it lengthens from **57 min at V_max_an = 0.20 to 75 min at the milder 0.15** — seed-invariant within each severity (IQR [57, 57] at 0.20 and [75, 75] at 0.15; Figure S2, Supplementary Methods S8.2) — because a slower acid-base decline pushes the pH < 7.15 crossing later while the DTH alarm stays early (~10 min throughout). The 57 min is therefore one point on a well-behaved envelope, not a universal value; we did not sweep harsher regimes (≥ 0.25), which enter the metabolic-runaway limitation (Section 5). Second, the evidence is associational (correlation plus relative alarm timing); establishing that the autonomic axis *causally leads* the acid crossing would require intervention experiments the testbed makes possible but that we have not yet run — lagged cross-correlation, and ablations that advance the acid-base trajectory while holding autonomic exhaustion fixed (and vice versa). We flag these as the natural next use of the instrument. These lead times should be read as internal model behavior under specified severity settings, not as expected clinical lead times.

*Methodological takeaway:* the physiological axis a biomarker correlates with, and the axis it usefully predicts, can differ; ground-truth labels let one separate the two instead of conflating them.

### 3.4 Sampling fidelity selects the biomarker

Applying the measurement model (Section 2.5) to a decompensation run, a deceleration-area biomarker and a beat-to-beat HRV biomarker (RMSSD) degrade in opposite ways as sampling rate falls. The deceleration area is **preserved to 100% of its beat-resolved value at every f_s down to 4 Hz** (107 269 bpm*·*s, unchanged from 4 to 250 Hz). RMSSD, by contrast, is corrupted by quantization jitter: at 4 Hz it is **inflated to ~808% of its native value** (138 ms vs a native ~17 ms), and it recovers to within ~10% only at ≥128 Hz (105% at 128 Hz, 101% at 250 Hz) (Figure 4). The inflation is the same overestimation-of-baseline-variability mechanism that, in vivo, made RMSSD fail to signal acidemia early at the clinical 4 Hz rate while succeeding at 1000 Hz [5, 4].

**Figure 4.**
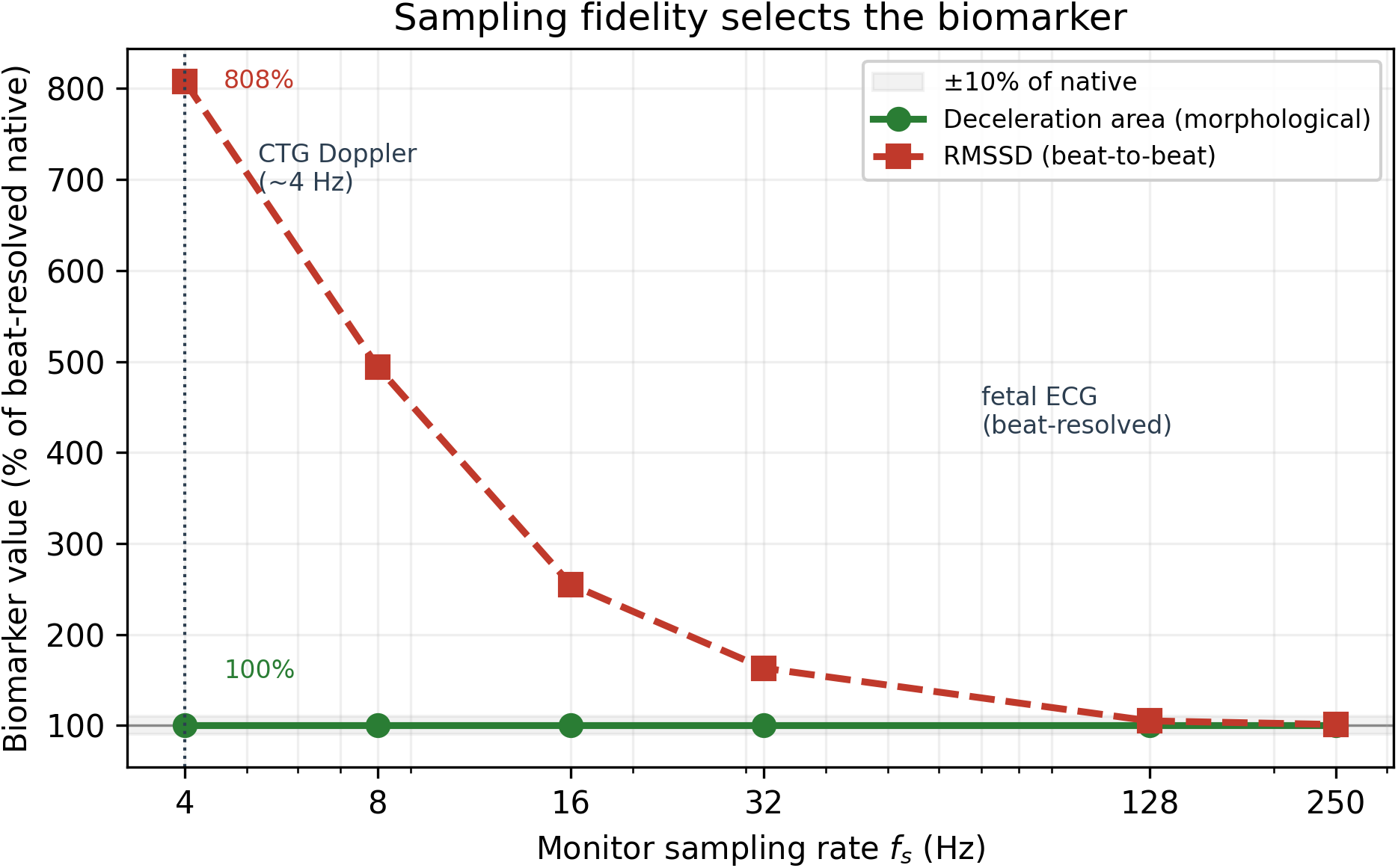
Sampling fidelity selects the biomarker. Deceleration-area (morphological) versus RMSSD (beat-to-beat) retention, as a percentage of the beat-resolved (native) value, across monitor sampling rates. The deceleration area is preserved at 100% of native down to CTG-Doppler 4 Hz, whereas RMSSD is inflated to 808% of native at 4 Hz by quantization jitter and recovers to within ~10% only at ≥ 128 Hz. Data: `output_fidelity/fidelity_sweep.csv` (`scripts/run_fidelity_sweep.py`).

The implication is a fidelity-matched selection rule: area-type biomarkers are deployable on CTG Doppler, whereas HRV biomarkers require fetal-ECG-grade sampling. This is the same design constraint that motivated the coarse-time-scale (2.5–8 s) features of Roux et al. (2021) [3], and the simulator reproduces it against known ground truth.

Two scoping caveats qualify this result. First, our measurement model isolates *timing quantization* alone; real Doppler CTG adds autocorrelative peak-detection uncertainty, signal dropout, internal smoothing/averaging, and proprietary preprocessing, all of which further degrade beat-to-beat metrics, so the 4 Hz result is an **idealized lower bound** on CTG corruption of RMSSD — a real monitor would do no better and generally worse — and the qualitative selection rule is therefore conservative. Second, the RMSSD failure at 4 Hz is not a property of this particular model: it follows from the arithmetic of quantizing sub-sample beat-to-beat differences onto a coarse grid, and is therefore **expected to generalize** to any beat-to-beat variability metric on any low-rate acquisition. The simulator’s contribution is to place this general effect against continuous ground truth and alongside a competing morphological biomarker on the same runs.

### 3.5 A negative result: rise-then-collapse is not reproducible from autonomic modulation here

We attempted to reproduce the published RMSSD “rise-then-collapse” trajectory by scaling phasic-HRV amplitude with vagal *activation*, so that early-hypoxia chemoreflex drive would amplify HRV before exhaustion collapses it. An A/B comparison over a delayed-exhaustion regime showed this **does not work**: the activation-scaled and fixed-amplitude RMSSD curves were nearly identical, both declining monotonically (Figure 5). In this model the vagal activation f_va never rises far enough above baseline, and exhaustion engages too quickly, for a rise to appear.

**Figure 5.**
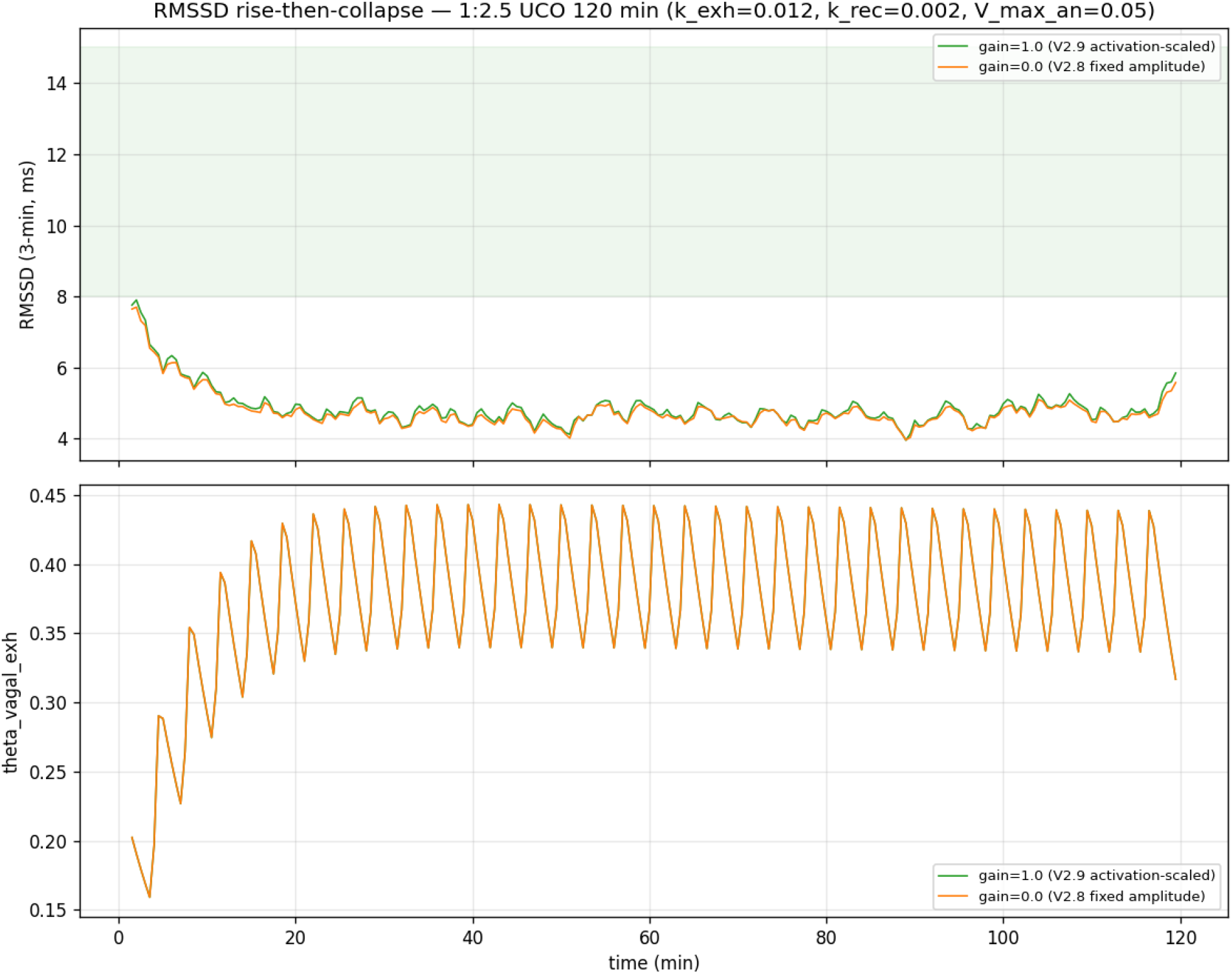
A negative result: rise-then-collapse is not reproducible from autonomic modulation here. RMSSD (3-min sliding window) over a 120-min 1:2.5 UCO run with phasic-HRV amplitude scaled by vagal activation (gain = 1.0) versus fixed amplitude (gain = 0); the vagal-exhaustion state *θ* is shown below. Both RMSSD curves decline monotonically — activation scaling does not reproduce the published rise-then-collapse — supporting that autonomic modulation alone, in this model class, is insufficient to reproduce the trajectory, and motivating an intrinsic-pacemaker hypothesis.

Stated carefully, the result is this: **autonomic modulation alone, in the present model class, is insufficient to reproduce the reported rise-then-collapse** trajectory (and, by extension, the nonlinear HRV-complexity changes seen in isolated-heart and chronic-hypoxia preparations [10] and in developmental-exposure cohorts [11]). An intrinsic-pacemaker contribution is one plausible explanation, but it is **not uniquely established** by these simulations: the failure could instead reflect missing or mis-parameterized autonomic physiology — chemoreflex timing, exhaustion dynamics, respiratory coupling, sympathetic re-engagement, or nonlinear state dependence — that a different autonomic-loop configuration might capture. What the negative result does do is *motivate*, rather than prove, rebuilding HRV bottom-up from a stochastic sinoatrial coupled-clock pacemaker with the autonomic loop as a modulatory overlay (future work; Section 4), and it identifies the phenomenon as one that a purely autonomic-overlay model should not be expected to generate.

## 4 Discussion

The demonstrations in Section 3 are of three deliberately distinct kinds. (i) **Artifact diagnosis** — beat-to-beat HRV amplitude can be a numerical artifact of the observation pipeline (3.1); this is the most secure kind of claim, because it is about the analysis, not the physiology. (ii) **Biomarker-design implications** — beat-to-beat variability requires an explicit stochastic driver in this model class (3.2), an HRV biomarker can correlate with one axis while usefully predicting an event on another (3.3), and biomarker information content depends quantifiably on acquisition fidelity (3.4); these hold for this model class and, in the case of 3.4, follow from acquisition mathematics that generalize beyond it. (iii) **Physiology hypothesis generation** — the failure to reproduce rise-then-collapse from autonomic modulation (3.5) motivates, but does not establish, an intrinsic-pacemaker account. None of the three is visible without ground-truth labels and the in-silico controls (noise-off, detector swap, fidelity quantization) that a mechanistic model uniquely permits.

### Model-based epistemology

What can an intervention inside an admittedly imperfect simulator legitimately establish? Robustly, it can establish *conditional, internal, comparative* facts: that a specific analysis choice fabricates structure (3.1), that a named mechanism is necessary for a behavior within the model (3.2), or that two biomarkers rank differently under a controlled degradation (3.4). It cannot, on its own, establish absolute physiological quantities, in-vivo timing, or which of several omitted mechanisms is responsible for a discrepancy. Our claims are pitched at the first level; where they touch the second (e.g. the ~57-min lead, the intrinsic-pacemaker account) we mark them as illustrative or hypothesis-generating.

### 4.1 What the model must — and must not — be relied on for

Because the model is calibration-grade rather than cohort-validated, we state its scope of validity explicitly. **The conclusions here require the model to reproduce only:** plausible deceleration morphology; the plausible ordering hypoxia → autonomic change → acid-base deterioration; the sampling-fidelity mathematics of beat-to-beat versus morphological metrics; and internally consistent, controllable ablations. Each of these is a qualitative or relational property that the model does reproduce, and each is what the corresponding claim actually rests on. **The model is *not* claimed to validate:** absolute quantitative fetal-HRV norms; exact in-vivo time-to-decompensation; transfer of specific numbers from fetal sheep to humans; or nonlinear pacemaker dynamics it does not explicitly represent. A reader should trust the conclusions exactly to the extent that they depend only on the first list — which is how they have been written.

This work is deliberately scoped as a methods-and-instrument contribution. The demonstrations use a single model of a single species (term fetal sheep), calibrated to plausible physiology but not fit to a specific experimental cohort; the antepartum growth scaling is validated to trend, not to absolute preterm norms. The value is as a controlled environment in which candidate biomarkers can be dissected against known ground truth before they are taken to real, unlabeled recordings. We are pursuing the intrinsic-pacemaker question (3.5) in a separate coupled-clock extension; it is reported here only as motivation, and its unpublished results are not presented.

### 4.2 Current status and roadmap

We state plainly where the instrument stands, because the distinction is central to what it can and cannot yet claim. The cardiovascular, metabolic/acid–base, and developmental (gestational growth-scaling) layers are **mechanistic**: their states evolve by integrated dynamics. The heart-rate-variability layer is the exception — it is presently a **distilled surrogate**: a per-beat autoregressive process (Section 2.1) that reproduces the *statistics* of intrinsic variability but does not *generate* them from pacemaker mechanism. A separate, mechanistic **coupled-clock pacemaker** — in which beat-to-beat variability emerges from sinoatrial channel and calcium-release noise — has been developed and evaluated in isolation against isolated-heart and chronic-hypoxia benchmarks; those results are outside the scope of the present paper and are not used to support any claim made here, and the pacemaker is **not yet embedded** in the labor model. The obstacle is well defined: the pacemaker requires a fixed-step stochastic (millisecond-resolution) integrator, whereas the labor model is a stiff, adaptively-integrated system spanning hours; coupling the two is a multi-timescale co-simulation problem, and the surrogate is the interim reduction that makes long labor runs tractable.

The defined next step is therefore to **replace the surrogate with the emergent pacemaker in the loop** — the step that would earn the HRV layer the same mechanistic standing as the rest of the model, and that is prerequisite to using the testbed for *discovery* of mechanism-dissociable HRV biomarkers (e.g. distinguishing hypoxic from inflammatory signatures), as opposed to interrogation of pre-specified ones. We stress that the findings reported here do **not** depend on this upgrade: the detector-artifact (3.1), injected-versus-emergent (3.2), fidelity-matched-selection (3.4), and lead-time (3.3) results are properties of the observation model and the acid-base coupling, and stand as reported. The emergent pacemaker *extends* the instrument’s reach; it does not revise these results.

## 5 Limitations

- **Calibration-grade, not validated**. Results reproduce plausible physiology; they are not fit to specific isolated-heart or chronic-hypoxia data. That validation is explicit future work.
- **Single model, single species** (term fetal sheep).
- **Metabolic runaway** past ~120–130 min in the aggressive anaerobic regime (`V_max_an = 0.2`): lactate exhausts the bicarbonate buffer and pH crashes unphysiologically; long runs require a milder setting. The model lacks bicarbonate regeneration / lactate-clearance saturation at extreme acidemia.
- **Spectral analysis is aliasing-limited** above ~1 Hz (mean beat rate ≈ 2 Hz).
- **Regime-bounded lead-time evidence**. The lead is invariant across five seeds and varies with severity in a well-behaved way (57 min at V_max_an = 0.20, 75 min at 0.15; Figures S1–S2), but the sweep is bounded to the physiologic regime — harsher severities (≥ 0.25) enter the metabolic-runaway regime and are not reported. The *ρ*(DTH, pH) value is seed-dependent (−0.34 to −0.78), and the causal-ordering interventions of Section 3.3 are not yet run.
- **Model-form uncertainty**. Several conclusions — especially the rise-then-collapse negative result (3.5) — may depend on which mechanisms are *omitted*, not merely on parameter settings; a different autonomic-loop structure could change them.
- **Detector- and analysis-pipeline dependence**. Some findings (3.1, 3.2) are statements about the observation and analysis model as much as the physiology, and may vary with specific implementation choices; we mitigate this by making the detector and noise models explicit and swappable, but do not claim invariance to them.
- **Intrinsic-HRV surrogate is phenomenological** (AR(2)), not a mechanistic sinoatrial model; claims about “intrinsic” origin are correspondingly provisional (Section 2.1).

## 6 Conclusion

**Fetal Twin** (the source-available `fetaltwin` engine) is a ground-truth-labeled mechanistic testbed for fetal physiological development, spanning antepartum growth and intrapartum stress. Its name’s digital-twin connotation is deliberate but bounded: it is a mechanistic, population-level twin used as a research instrument, not a validated, patient-specific clinical digital twin or predictor. Its primary value is as a controlled environment in which candidate fetal-monitoring biomarkers can be dissected against known latent labels. Demonstrated on a fetal-sheep-derived autonomic-loop configuration, it shows that beat-to-beat HRV amplitude can be a numerical artifact of the observation pipeline; that beat-to-beat variability in this model class requires an explicit stochastic driver rather than arising from the deterministic loop; and that biomarker information content depends on acquisition fidelity in a way that follows from quantization mathematics — a fidelity-matched selection rule. A negative result — the non-reproducibility of the published rise-then-collapse from autonomic modulation — motivates, without proving, an intrinsic-pacemaker account. The instrument and these demonstrations are offered to the fetal-monitoring and physiological-modeling communities; the engine and the configurations reproducing every figure are released for noncommercial research.

## 7 Code and data availability

### What is available

The simulation engine (the full ODE model and both right-hand-side backends), the scenario configurations, and the analysis scripts that generate the figures in this paper (with one exception noted below) are released as the Fetal Twin engine at https://github.com/martinfrasch/fetaltwin (project home https://fetaltwin.org) under the PolyForm Noncommercial License 1.0.0, archived at Zenodo (DOI: 10.5281/zenodo.21158927). The release also includes a browser-based interactive demonstration (a Streamlit application) that runs the engine, builds umbilical-cord-occlusion scenarios, replays a simulation as a scrolling fetal-monitor (cardiotocography) strip, and browses the model equations — lowering the barrier to exploring the testbed. It is a research/demonstration front-end, not a medical device. Academic and other noncommercial users may run the complete engine and reproduce the reported experiments — including the antepartum growth-scaling baselines and the intrapartum demonstrations (the beat-detector artifact, the injected-versus-emergent spectral test, the sampling-fidelity result, and the rise-then-collapse negative result) — from the released configurations and seeds. **One exception:** the Distance-to-Healthy figures (Figures 3, S1, S2) rely on an external HRV-feature library for their beat-to-beat feature extraction, invoked by (but not bundled with) the released scripts; those specific figures are therefore not reproducible from this release alone. Separately, no result in this paper depends on any deliberately withheld production asset (below). A Supplementary Methods appendix (scenario configs, seed handling, exact parameter deltas from Wang 2015, backend equivalence, detector comparison, fidelity-sweep procedure, and the predictor alarm criterion) accompanies the release and this submission. The installable Python package currently retains the internal module name `labor_sim`; the public repository and project name are `fetaltwin`.

### What is not part of the release, and why

Production-calibrated multi-axis parameter corpora and any trained-predictor weights developed for commercial use are *not* included; these are maintained as trade secrets by JoyBeat Medical, Inc. They are **not required to reproduce any figure or claim in this paper** — the results here use only the illustrative/default configurations that ship with the open engine. Methods implemented in the software are the subject of a pending U.S. patent application assigned to JoyBeat Medical, Inc.; commercial use requires a separate license. This open-core arrangement is intended to make the instrument fully usable and reproducible for research while reserving commercial rights.

## 10 Acknowledgements

This work is built on the combined fetal-sheep umbilical-cord-occlusion model of Wang et al. (2015) [1]. The author was a member of that original group and gratefully thanks his co-authors — Q. Wang, N. Gold, C. L. Herry, M. G. Ross, and B. S. Richardson — for laying the mathematical and physiological foundation on which the present testbed extends, and acknowledges the funding bodies that supported that original work. Implementation and manuscript drafting were assisted by a large language model (Anthropic Claude) under the author’s direction and review. The author is responsible for all content.

